# Comparison of the bioactivity of gold, silver and copper oxide nanoparticles derived from *Aloe africana* leaf extract and *Magnetospirillum magnetotacticum*

**DOI:** 10.64898/2026.02.12.705669

**Authors:** Khanyisile Ngcongco, Anushka Govindsamy, Nirasha Nundkumar, Karen Pillay

## Abstract

Metallic nanoparticles have emerged as novel therapeutic agents due to their distinctive physicochemical properties and broad-spectrum activity, with applications in antimicrobial therapy, drug delivery and bioremediation. Conventional methods for metallic nanoparticle synthesis often utilize toxic chemicals and energy intensive processes that are expensive. Green synthesis offers a sustainable and cost-effective alternative by using biomolecules from plants and microorganisms. In this study, gold (AuNPs), silver (AgNPs), and copper oxide (CuO NPs) nanoparticles were biosynthesized using leaf extracts of *Aloe africana Mill*., a South African medicinal plant rich in phytochemicals, and the magnetotactic bacterium *Magnetospirillum magnetotacticum* that naturally produces intracellular nanoparticles. GC-MS analysis revealed 13 known phytochemicals in the *A. africana* extract including esters, terpenoids, monoglycerides, and fatty acids which served as reducing and capping agents for nanoparticle synthesis. *A. africana*-derived AgNPs were spherical (11-30 nm) in shape, capped with dihydrosqualene, a known antibacterial compound; and was found to display activity against Gram-negative (*Escherichia coli* and *Pseudomonas aeruginosa*) and Gram-positive (*Enterococcus faecalis* and *Staphylococcus aureus*) bacteria. These AgNPs however exhibited cytotoxicity to HEK293 and HeLa cell lines. *A. africana* AuNPs (17-62 nm) displayed diverse morphologies and CuO NPs (55-115 nm) were irregular shaped, and both nanoparticles exhibited limited antibacterial activity and low cytotoxicity. *M. magnetotacticum*-derived AuNPs (12-21 nm) and AgNPs (51-126 nm) were spherical, with the CuO NPs (42-66 nm) having irregular shapes. Except for *A. africana*-derived AgNPs, all other metallic nanoparticles displayed poor antibacterial activity. These findings are novel and highlight a dual-function green synthesis platform where *A. africana* phytochemicals contribute to both nanoparticle synthesis and bioactivity, positioning *A. africana* AgNPs as promising antibacterial agents.

## 1. Introduction

The emergence of bacteria that are resistant to antibiotics has highlighted the need for alternatives to currently available antibiotics (Keshari *et al*., 2020). Metallic nanoparticles have been found to display antibacterial activity against Gram-negative and Gram-positive bacteria, leading to their investigation as potential antibacterial agents (Wang *et al*., 2017). These nanoscale structures possess remarkable physicochemical properties and show promise for use in a wide variety of applications in biomedicine, biotechnology, environmental science, and engineering (Akbarzadeh *et al*., 2012). Metallic nanoparticles are synthesised with metal precursors using conventional synthesis methods i.e. chemical and physical synthetic strategies (Rambau *et al*., 2024). However, these synthesis methods often involve the use of toxic reagents, incur high production costs, and generate environmental pollution, which present significant drawbacks (Satyavani *et al*., 2011; Chikdu *et al*., 2015). Therefore, in response to these challenges, there is a need for alternative nanoparticle synthesis strategies that are efficient and not harmful to the environment. Green synthesis methods have opened a door to innovative nanoparticle production using cost effective and eco-friendly processes.

Green synthesis strategies use biological organisms such as plants, bacteria, yeast, fungi, or algae to mediate nanoparticle synthesis (Nadaroglu *et al*., 2017). Plants are promising candidates for nanoparticle synthesis due to their ability to detox and reduce the accumulation of metals by altering the chemical composition of metals making them non-toxic and thus producing nanoparticles as a by-product (Nadaroglu *et al*., 2017). A wide variety of plant-derived biomolecules including organic acids, proteins, enzymes, phenolic compounds, vitamins, alkaloids, flavonoids, terpenoids, polysaccharides, amines, and pigments play crucial roles in the reduction and stabilization of nanoparticles during synthesis (Nadaroglu *et al*., 2017; Ijaz *et al*., 2020; Akintelu *et al*., 2021). These plant biomolecules can be extracted from various plant parts such as the leaves, stems, roots, fruit, bark, flowers, seeds, and buds and used for synthesising nanoparticles (Ijaz *et al*., 2020; Yulizar *et al*., 2020). While numerous plant species have been explored, there is growing interest in the use of medicinal plants in green nanotechnology. Medicinal plants are an invaluable source of natural bioactive compounds (Xulu *et al*., 2022; Puja *et al*., 2023). Approximately 80% of the global population continues to rely on medicinal plants for their primary healthcare and for the development of medicines (Xulu *et al*., 2022). *Aloe africana* commonly known as the African aloe belongs to the *Aloe* genus comprising over 430 species (Nalimu *et al*., 2021). Aloe plants contain a wide variety of bioactive compounds including flavonoids, terpenoids, lectins, fatty acids, anthraquinones, tannins, sterols, enzyme, vitamins, and saccharides (Hęś *et al*., 2019). These bioactive compounds are associated with diverse biological activities which include antifungal, antibacterial, antiviral, anti-inflammatory, antimicrobial, laxative, immunomodulating and anticancer effects (Guo and Mei, 2016).

In addition to plants, bacteria are also suitable candidates for nanoparticle synthesis. Bacteria offer several advantages, including rapid growth, cost effectiveness, controlled culturing conditions and the ability to manipulate their growth conditions and environment (Pantidos and Horsfall, 2014; Nadaroglu *et al*., 2017). These advantages can be exploited for nanoparticle production. Certain Gram-negative and Gram-positive bacterial species have the ability to mitigate metal and heavy metal toxicity, allowing them to facilitate nanoparticle formation under these toxic conditions (Nadaroglu *et al*., 2017). *Magnetospirillum magnetotacticum* (MTB) is a bacterial species that belongs to a group of Gram-negative bacteria known as magnetotactic bacteria that can biomineralize iron ions and convert them into magnetic nanoparticles such as magnetite (Fe_3_O_4_) or greigite (Fe_3_S_4_) in vesicle-like structures enveloped by a phospholipid bilayer membrane, termed magnetosomes (Scheffel *et al*., 2008; Ge *et al*., 2011; Lin *et al*., 2020).

Various metallic nanoparticles such as silver, gold, copper, selenium, nickel, zinc oxide, titanium oxide, and iron oxide have been studied for their antimicrobial efficacy due to their unique physicochemical properties (Ozdal and Gurkok, 2022). The antibacterial activity of these nanoparticles is primarily due to their ability to interact electrostatically with the bacterial cell membranes, leading to the formation of pores or pits which compromise the membrane integrity (Dong *et al*., 2019; Yu *et al*., 2020). This disruption causes membrane rupture and subsequent cell shrinkage (Yu *et al*., 2020). Following membrane interaction, nanoparticles enter the cell via endocytosis, and the nanoparticles are localized in lysosomes where oxidation facilitate the release of metal ions (Ngcongco *et al*., 2023). These metal ions subsequently interact with critical cell components and biomolecules including lipids, proteins, nucleic acids, ribosomes, and mitochondria, inducing cell damage (Zhang *et al*., 2016; Sharifi-Rad *et al*., 2020). Metal ions can induce DNA damage by causing strand breaks, disrupting the double helical structure, and causing mutations in the DNA genes responsible for repairing DNA, thereby inhibiting DNA replication and triggering apoptosis (Zhang *et al*., 2016). Additionally, mitochondrial dysfunction disrupts the electron transport chain and promotes apoptosis (Beyth *et al*., 2015). Thus, nanoparticles have multiple antibacterial mechanisms compared to traditional antibiotics making them promising alternatives as antibacterial agents.

Despite the growing interest in green synthesis, no prior studies have investigated *A. africana* and *M. magnetotacticum* for comparative biosynthesis of gold, silver and copper oxide nanoparticles and for their bioactivity. The nanoparticles produced by the two green strategies were evaluated for broad-spectrum antibacterial activity against Gram-negative and Gram-positive pathogens and their cytotoxic effect against normal and cancerous human cell lines, thereby delivering insights into sustainable antimicrobial nanotechnology from two distinct biological sources. Through GC-MS, phytochemical profiling was used to identify specific reducing and capping agents in *A. africana* extracts and their retention on nanoparticle surfaces, thereby providing invaluable data on the therapeutic potential of this aloe species.

## 2. Materials and methods

### 2.1. Materials

*Aloe africana* plants were obtained from a local South African nursery specialising in the retail of indigenous species. All chemicals, solvents and media used in this study were of analytical grade and purchased from Merck (Pty) Ltd, South Africa, unless stated otherwise. Antibiotics were purchased from Sigma Aldrich, Germany. *M. magnetotacticum* strain MS-1 was purchased from Deutsche Sammlung von Mikro-organismen und Zellkulturen (DSMZ, Germany). The following strains were from the American Type Culture Collection (ATCC): *Escherichia coli* (ATCC 35218), *Enterococcus faecalis* (ATCC 5129), *Pseudomonas aeruginosa* (ATCC 27853) and *Staphylococcus aureus* (ATCC 43300).

### 2.1. Extract preparation

The extraction of *Aloe africana* phytochemicals was carried out using a modified method adapted from Gupta and Tejavath (2019). Fresh *A. africana* leaves were washed with distilled water to remove impurities. The leaves were cut into small pieces, suspended in Milli Q water, crushed and boiled in an Erlenmeyer flask, then allowed to cool to room temperature, filtered and stored in the dark at -20°C.

### 2.2. Nanoparticle synthesis from *A. africana* leaf extract

The synthesis of nanoparticles using the aqueous *A. africana* leaf extracts was done using a modified method adapted from Gupta and Tejavath (2019) and Arsène *et al*. (2023). *A. africana* extract was reacted with metal salt solution followed by incubation. Nanoparticle production was identified by a colour change, for gold nanoparticles (AuNPs) – from a yellow to a violet or reddish-brown precipitate, for silver nanoparticles (AgNPs) – from clear to a brown precipitate, and for copper oxide nanoparticles (CuO NPs) – from a greenish blue to a brown precipitate. The formed nanoparticles were pelleted, washed, and dried in an oven. The dry yield of the nanoparticles was determined, followed by resuspension in distilled water and storage in the dark at -20°C.

### 2.3. MTB stock preparation

Chemically defined media (CDM) was prepared for growing the bacteria *Magnetospirillum magnetotacticum* MS-1, and its composition is listed in Table 1. The mixture was then autoclaved at 120°C for 15 minutes. *Magnetospirillum magnetotacticum* MS-1 (10% vol/vol) was inoculated into the CDM and the cultures were incubated at 30°C for 24 hours. Optical density (OD) readings of the samples were carried out at an absorbance of 565 nm and were obtained before and after incubation to monitor bacterial growth.

**Table 1:** Chemically defined media composition (1 L)

| Solution | Components | Quantity |
| --- | --- | --- |
| **ATM4S solution**<br>*Make up with deionized water (18 mΩ) to 295 ml* | Ammonium sulphate (NH<sub>4</sub>)<sub>2</sub>SO<sub>4</sub> | 4 g |
|  | Sodium dihydrogen phosphate (NaH<sub>2</sub>PO<sub>4</sub>·H<sub>2</sub>O) | 5.2 g |
|  | Di-potassium hydrogen phosphate (K<sub>2</sub>HPO<sub>4</sub>) | 12.5 g |
|  | Tri-sodium citrate (Na<sub>3</sub>-citrate·2 H<sub>2</sub>O) | 0.66 g |
|  | NaNO<sub>3</sub> | 0.12 g |
| **Glucose solution**<br>*Make up with deionized water (18 mΩ) to 250 ml* | Glucose Monohydrate | 1 g |
|  | Magnesium sulphate heptahydrate | 0.050 g |
| **Yeast extract solution**<br>*Make up with deionized water (18 mΩ) to 150 ml* | Yeast Extract | 10 g |
| **Trace metal stock solution**<br>*Make up with deionized water (18 mΩ) to 1 L* | Na-EDTA | 20.1 g/L |
|  | FeCl<sub>3</sub>·6 H<sub>2</sub>O | 16.7 g/L |
|  | ZnCl<sub>2</sub> | 0.13 g/L |
|  | CoCl<sub>2</sub>·6 H<sub>2</sub>O | 0.18 g/L |
|  | NaMoO<sub>4</sub>·2 H<sub>2</sub>O | 0.13 g/L |
|  | CaCl<sub>2</sub> | 0.50 g/L |
|  | CuSO<sub>4</sub>·5 H<sub>2</sub>O | 0.16 g/L |
|  | H<sub>3</sub>BO<sub>3</sub> | 0.28 g/L |
|  | MnCl<sub>2</sub>·2 H<sub>2</sub>O | 0.11 g/L |

Individual components of this media were prepared as described in Table 1.
Thereafter, 295 ml of ATM4S solution, 250 ml of glucose solution, 150 ml of yeast extract solution, and 2 ml of trace metal stock solution were combined and were made up to 1 L with distilled water and then the pH was adjusted to 6.8.

### 2.4. MTB nanoparticle synthesis

The synthesis of nanoparticles using *M. magnetotacticum* was conducted with slight modification to the method described by Murei *et al*. (2021). A 10 mM metal salt stock solution of the metals silver, gold, and copper was made up in 50 ml tubes by calculating the dry mass of the metals’ chloride form that would be required to yield a stock of accurate concentration. The metal chloride was then mixed with autoclaved distilled water, vortexed and stored in the fridge. Metal stock solutions (4 ml) were added to 496 ml of chemically defined media without yeast extract, containing 0.5 mg ISOGRO^®^. *M. magnetotacticum* was inoculated into the media containing the respective metals at a starting OD range of 0.5 to 0.6. The tubes were wrapped in foil to prevent photodegradation of the metals in the media. The tubes were then incubated at 30°C for 4 days. Following the incubation period, the cultures were centrifuged at 10 000 rpm for 30 minutes at 4°C to pellet the cells. The pellet was resuspended in a 5 ml volume of 20 mM HEPES and 20 mM NaCl buffer (pH 7.4). The suspension was sonicated for 30 minutes at 20°C to break open the cells thereby releasing the nanoparticles. The suspension was then centrifuged at 4 000 rpm for 30 minutes at 4°C. The supernatant was collected and added to pre-weighed tubes, then dried in the oven set at 40-50°C. The weight of the tubes was taken once the nanoparticles were dried to determine nanoparticle dry yield.

### 2.5. UV-Vis spectral analysis

The synthesis of gold (AuNPs), silver (AgNPs), and copper oxide (CuO NPs) nanoparticles by reduction of aqueous Au³□, Ag□, and Cu²□ precursors was confirmed using UV-vis spectrophotometric monitoring. This was done by diluting a small aliquot of sample in diluted water and performing UV-Vis spectral analysis at a wavelength range of 200 – 800 nm. The expected absorbance maxima wavelength range of the AuNPs, AgNPs and CuO NPs nanoparticles are 500 – 600 nm, 400 – 500 nm and 250 – 350 nm, respectively.

### 2.6. TEM-EDX analysis

High-resolution transmission electron microscopy (HRTEM) analysis was carried out using a JEOL JEM-2100 electron microscope (JEOL Ltd., Korea). The nanoparticle samples were transferred onto carbon-coated copper grids (Formvar film), while nickel grids were specifically employed for the characterization of copper oxide nanoparticles. The grids were allowed to air-dry completely prior to imaging. HRTEM imaging was conducted at multiple magnifications to elucidate the morphological and structural features of the nanoparticles. Additionally, elemental composition and distribution were determined via energy-dispersive X-ray spectroscopy (EDX) using an Oxford X-Max 18 mm silicon drift detector (SDD) system (Oxford Instruments, UK) integrated with the HRTEM.

### 2.7. Dynamic light scattering (DLS) analysis

To determine the size distribution profile and surface charge of the nanoparticles in an aqueous medium, the zeta size and zeta potential was measured using a Malvern pan analytical, Zetasizer nano ZS instrument operating on software version 8.02 (United Kingdom).

### 2.8. Gas chromatography-mass spectrometry (GC-MS)

Leaf extracts and plant extract-coated nanoparticles were suspended in analytical-grade methanol and sonicated (10 min at 20°C) to release surface-bound phytochemicals for volatilization and GC-MS analysis. The analyses of the extracts and nanoparticles were performed using a GCMS-QP 2010SE instrument (Shimadzu, Japan) on a Zebron ZB-5MSplus, 30 m x 0.25 mm (0.25 µm film) column. The carrier gas, helium, was of ultra-high purity grade (Afrox, Gauteng, South Africa) and was set at a flow rate of 1.4 ml min^−1^ in the constant flow mode. The SPME fibre was desorbed for 5 min in a SPME inlet liner (Supelco, Sigma-Aldrich (Pty) Ltd. Kempton Park, South Africa) of a GC inlet at 225 °C. The GC inlet was operated in the split mode (100:1 split ratio). The GC oven temperature programme was 40 °C (0.9 min) at 8 °C min^−1^–280 °C (5 min). The GC run time was 35.9 min. The MS transfer line temperature was set at 280 °C and the ion source temperature was set at 200 °C. The electron energy was 70 eV in the electron impact ionization mode (EI +), the data acquisition rate was 10 spectra s^−1^, the mass acquisition range was 40–500 Daltons, and the detector voltage was set at 1650 V. The data were captured using the GC-MS workstation solution software (Shimadzu, Japan).

While FTIR commonly characterizes surface functional groups on green-synthesized nanoparticles, GC-MS enables detailed identification and quantification of desorbed volatile/semivolatile compounds providing complementary molecular insights.

### 2.9. Identification of compounds

Compounds were identified by comparing mass spectra (including molecular ions, fragments, and base structures) to the NIST library (National Institute of Standards and Technology, USA) standards. The retention time was matched, and the relative amounts of individual compounds were shown as the percentage peak areas relative to the total peak area. Only peaks with similarity index of ≥ 70% (as recommended by Rahim *et al*. (2017)) when matched against the NIST libraries were selected for identification. These peaks were further validated with Chemspider data and previously reported studies on phytochemical profiles of *Aloe* species and other plant leaf extracts (Table S1). Instrument related artifacts and potential contaminants were excluded from the analysis.

### 2.10. Antibacterial assay

The antibacterial activity assay was carried out using a modified method described by Moodley *et al*. (2018). Varying concentrations (6.25 - 200 µg/ml) of *A. africana* leaf extracts and AuNPs, AgNPs and CuO NPs synthesised using *A. africana* leaf extracts or MTB were tested for bioactivity against *Escherichia coli* (ATCC 35218), *Enterococcus faecalis* (ATCC 5129), *Pseudomonas aeruginosa* (ATCC 27853) and *Staphylococcus aureus* (ATCC 43300) using the iodonitrotetrazolium chloride (INT) assay. Serial two-fold concentrations of AuNPs, AgNPs and CuONPs were prepared in sterile 96-well plates over a range of 6.25 - 200 µg/ml. The first row of the plates was reserved for negative or uninhibited growth, containing bacteria only. The wells were then inoculated with diluted overnight bacterial culture adjusted to 0.5 McFarland standard and incubated at 30°C for 24 hours. Following the incubation period, 0.2 mg/ml of freshly prepared iodonitrotetrazolium chloride (INT) was added to the wells and the plates were incubated in the dark at 30°C for 4 hours. Neomycin served as a positive control. The absorbance was determined at 490 nm using a multimodal plate reader (Biotek Synergy HT, USA) with Gen 5 software (Biotek Synergy HT, USA Ver 2.01.14). The experimental design included two biological replicates, and in each instance triplicate samples were used.

### 2.11. Cytotoxicity Assay

The MTT assay was carried out using a modified method described by Naidoo *et al*. (2022) to determine cytotoxicity. Two human cell lines viz. embryonic kidney (HEK293) and cervical carcinoma (HeLa) were maintained in Eagle’s minimal essential medium (EMEM) containing 10% (v/v) gamma-irradiated FBS and 1% antibiotics (100 U/mL penicillin, 100 µg/mL streptomycin) at 37°C and 5% CO_2_, in a HEPA Class 100 Steri-Cult CO_2_ incubator (*Thermo-Electron Corporation,* Waltham, Massachusetts, USA). For the MTT assay, the cells were trypsinized and seeded into 96-well plates at a seeding density of approximately 3 x 10^4^ cells per well and incubated at 37°C for 24 h. The medium was replaced with fresh medium, thereafter, varying concentrations (31.25 – 1000 µg/mL) of *A. africana* extracts and *A. africana* nanoparticles were added to the wells followed by incubation at 37°C for 48 h. A negative control containing only cells was set up and this was set at 100% cell survival/viability. The assays were performed in triplicate. Following the incubation period, the spent medium was removed and 100 μL fresh medium and 10 μL MTT reagent (5 mg/mL in PBS) were added to each well. The cells were incubated for a further 4 h at 37°C. Thereafter, the MTT-medium was removed and replaced with 100 μL dimethylsulphoxide (DMSO) to solubilize the formazan crystals. The absorbance was read at 540 nm using a *Mindray* MR-96A microplate reader (*Vacutec*, Hamburg, Germany).

### 2.12. Statistical analysis

All data were statistically analysed using GraphPad InStat version 3.10 for Windows (GraphPad Software, San Diego California USA). Analysis of variance (*ANOVA*) was used, and differences were considered statistically significant at p < 0.05.

## 3. Results

### 3.1. UV-vis spectral analysis

The ultraviolet-visible (UV-vis) absorption spectra presented in Figure 1 confirms the successful biosynthesis of gold (AuNPs), silver (AgNPs), and copper oxide (CuO NPs) nanoparticles mediated by *A. africana* leaf extracts. Characteristic surface plasmon resonance (SPR) bands were observed at distinct wavelength maxima of approximately 590 nm for AuNPs, 440 nm for AgNPs, and 300 nm for CuO NPs.

**Figure 1:**
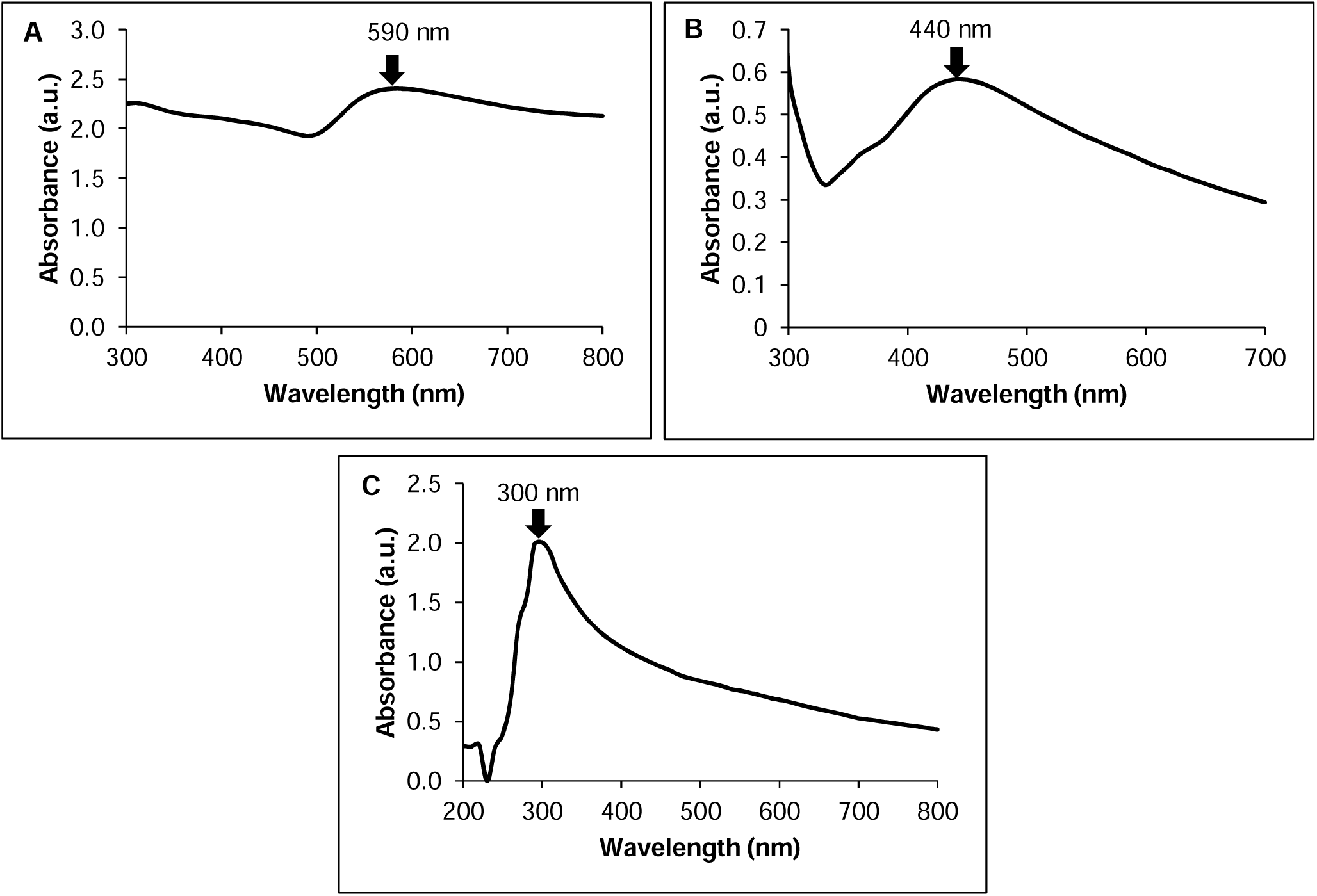
UV-vis spectra of nanoparticles (A) AuNPs, (B) AgNPs and (C) CuO NPs synthesized from *A. africana* leaf extracts.

UV-vis spectral analysis of bacterially synthesised nanoparticles yielded no defining peaks. This absence of defining peaks, rather than indicating failed synthesis, aligns with literature reports where TEM primarily provides confirmation of nanoparticle synthesis using these species (Murei *et al*., 2021; Sancho *et al*., 2023).

### 3.2. TEM-EDX analysis

The gold, silver and copper oxide nanoparticles synthesised from *A. africana* leaf extract had yields of 1.36 mg/ml, 1.19 mg/ml and 0.18 mg/ml, respectively, while those synthesised from 50 ml of *M. magnetotacticum* liquid culture had lower yields of 0.46 mg/ml, 0.71 mg/ml and 0.50 mg/ml, respectively. Transmission electron microscopy (TEM) and energy-dispersive X-ray spectroscopy (EDX) was conducted to determine the morphological features and elemental composition of the *A. africana* and *M. magnetotacticum* nanoparticles. TEM imaging of the *A. africana* derived AuNPs indicated spherical, hexagonal, triangular rod-like and diamond shaped nanoparticles with a size range of 13-47 nm (Figure 2A). EDX analysis confirmed the presence of Au as seen in Figure 2B. The *A. africana* AgNPs were more uniformly shaped as they were mostly spherical and oval shaped and slightly smaller with a narrow size distribution of 11-30 nm as seen in Figure 2C. The presence of Ag was confirmed by EDX analysis (Figure 2D). The TEM micrograph of CuO NPs (Figure 2E) displayed irregular shaped larger particles with a size distribution of 55-115 nm, and the EDX spectrum confirmed the presence of copper (Figure 2F).

**Figure 2:**
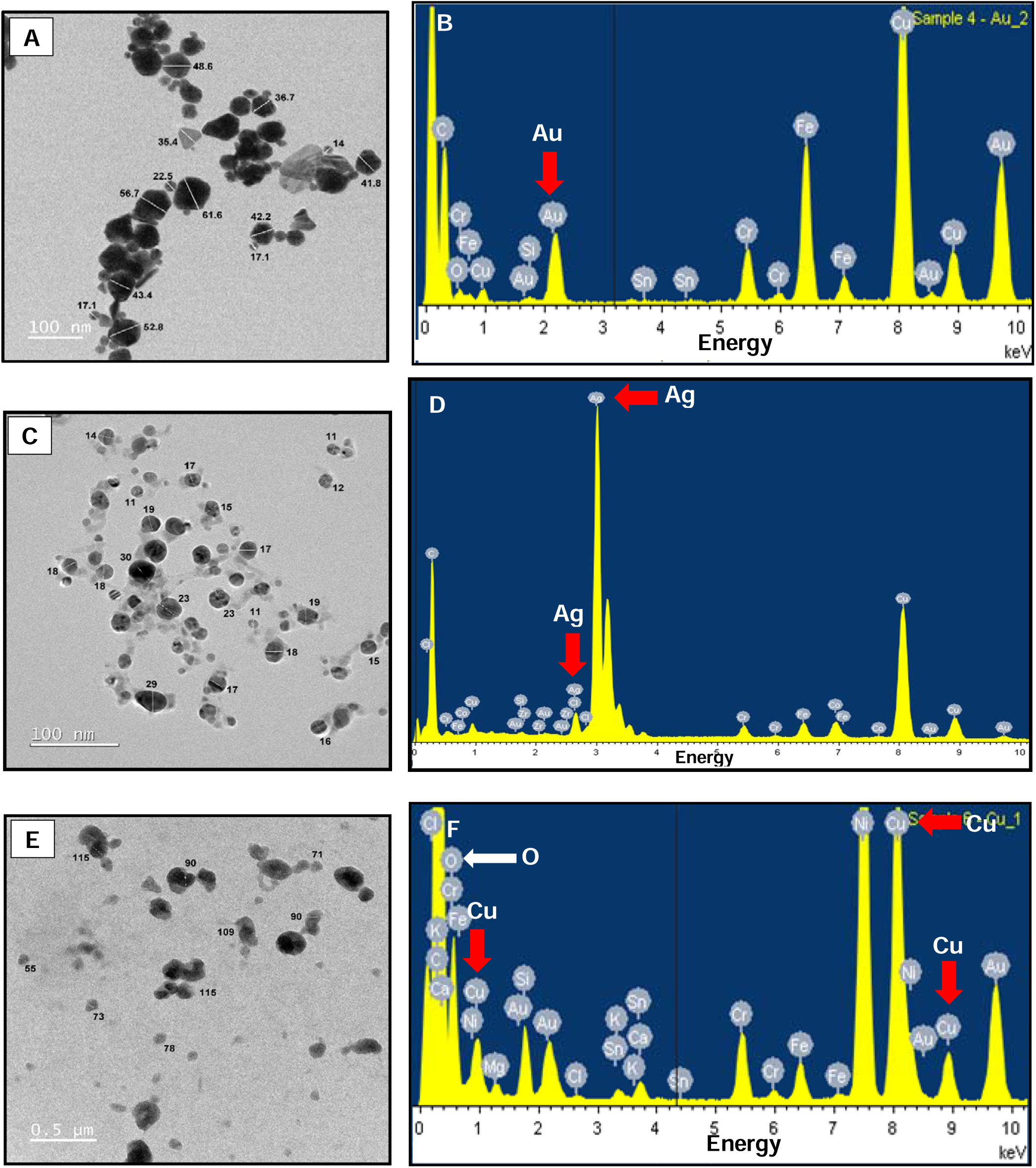
TEM micrographs showing (A) AuNPs, (C) AgNPs and (E) CuO NPs formed from *A. africana* leaf extract. Size ranges of AuNPs, AgNPs and CuO NPs nanoparticles are 17.1-61.6 nm, 11-30 nm and 55-115 nm, respectively. The EDX profiles of the (B) AuNPs, (D) AgNPs and (F) CuO NPs confirmed peaks of the respective metals.

TEM imaging of the *M. magnetotacticum* synthesized AuNPs and AgNPs indicated that the nanoparticles were spherical in shape. The AuNPs had a narrow size range of 12-21 nm (Figure 3A). EDX analysis confirmed the presence of Au as seen in Figure 3B. AgNPs had a size distribution of 51-126 nm as seen in Figure 3C and the presence of Ag was confirmed by EDX analysis (Figure 3D). The TEM micrograph of CuO NPs (Figure 3E) displayed irregular shaped particles with a size distribution of 42-66 nm, and the EDX spectrum confirmed the presence of copper (Figure 3F). Carbon and nickel are observed on the spectra and these elements are due to the grid used, background or instrumental artefacts. Hence, blank measurements were conducted, whereby EDX analysis was performed on grids without sample to establish background spectrum baseline. The blank measurements contained elemental peaks for chromium (Cr), iron (Fe), silicon (Si), copper (Cu) for copper grids and nickel (Ni) for nickel grids. These blank measurements were used for distinguishing between real elemental peaks due to the sample and background noise.

**Figure 3:**
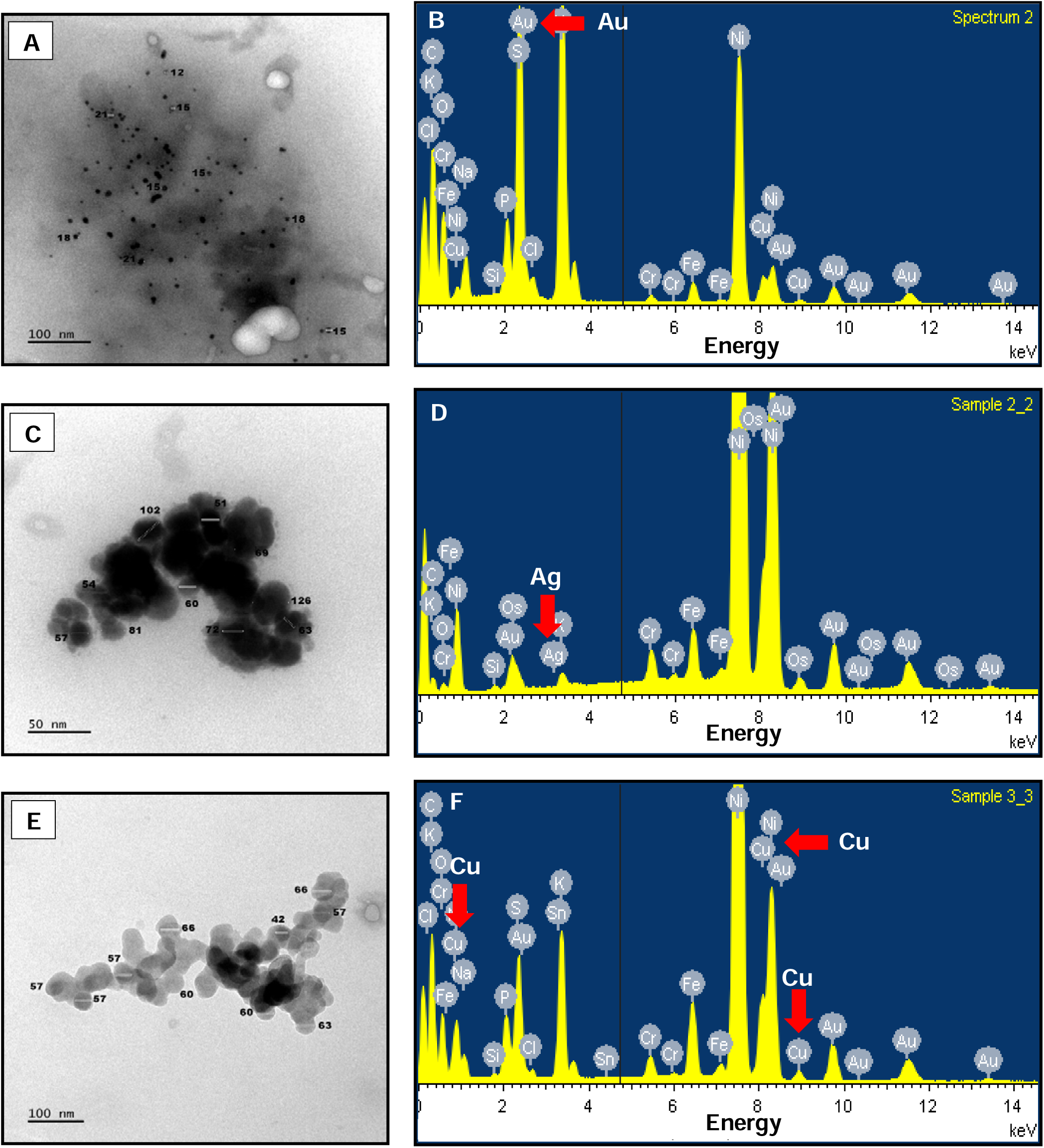
TEM micrographs of (A) AuNPs, (C) AgNPs and (E) CuO NPs synthesised by *M. magnetotacticum*. Size ranges of AuNPs, AgNPs and CuO NPs nanoparticles are 12-21 nm, 51-126 nm and 42-66 nm, respectively. The EDX profiles of the (B) AuNPs, (D) AgNPs and (F) CuO NPs confirmed peaks for the respective metals.

### 3.3. Dynamic light scattering (DLS) analysis

Dynamic light scattering (DLS) analysis was used to determine the dispersity and colloidal stability of the biosynthesized nanoparticles. The DLS measurements yielded the hydrodynamic diameter, particle size distribution, and zeta potential values, which are summarized in Table 2.

**Table 2:** Polydispersity index and zeta potential of *A. africana* and *M. magnetotacticum* nanoparticles.

| Nanoparticles | Polydispersity Index (PDI) | Zeta Potential (mV) |
| --- | --- | --- |
| *A. africana* AuNPs | 0.267 | -36.7 |
| *A. africana* AgNPs | 0.383 | -23.1 |
| *A. africana* CuO NPs | 0.336 | -16.7 |
| *M. magnetotacticum* AuNPs | 0.255 | -19.0 |
| *M. magnetotacticum* AgNPs | 0.402 | -29.7 |
| *M. magnetotacticum* CuO NPs | 0.302 | -28.4 |

PDI indication: values <0.1 indicate high monodispersity, values between 0.1-0.4 indicate moderate dispersity and values >0.4 indicate high polydispersity. Zeta potential indication: ± to 0-10 mV represent highly unstable, ± 10–20 mV represent relatively stable, ± 20–30 mV represent moderately stable, and $> \pm$ 30 mV represent highly stable nanoparticles.

The results demonstrate that nanoparticles synthesised using both *A. africana* and *M. magnetotacticum* exhibit moderate dispersion. Among these, the *M. magnetotacticum*-mediated silver nanoparticles displayed the smallest hydrodynamic size, whereas the copper nanoparticles synthesised using *A. africana* presented the largest hydrodynamic size. The zeta potential revealed that *A. africana*-mediated AuNPs exhibited high colloidal stability, the AgNPs showed moderate stability, and the CuO NPs were relatively stable. In contrast, *M. magnetotacticum*-mediated AuNPs demonstrated relative stability, while their AgNPs and CuO NPs were moderately stable.

### 3.4. Gas chromatography-mass spectrometry (GC-MS) analysis

The constituents present in the aqueous leaf extracts of *A. africana* and attached to *A. africana*-derived AuNPs, AgNPs and CuO NPs were identified by GC-MS analysis. A total of thirteen compounds with their retention time, molecular formula, molecular weight and percentage composition in the leaf extract are presented in Table 3. Compounds involved in nanoparticle synthesis are shown in Table 4, while those coating nanoparticle surfaces are presented in Table 5. Structures and biological activities of major compounds are detailed in Table 6.

**Table 3:** Compounds present in *A. africana* leaf extract.

| Retention Time (Min) | Name of compound | Molecular formula | Molecular weight (g/mol) | Peak Area % |
| --- | --- | --- | --- | --- |
| 5.723 | 4,8-Dimethylundecane | C<sub>13</sub>H<sub>28</sub> | 184 | 2.37 |
| 6.276 | 2-Bromononane | C<sub>9</sub>H<sub>19</sub>Br | 206 | 1.36 |
| 6.495 | E-2-Hexadecacen-1-ol | C<sub>16</sub>H<sub>32</sub>O | 240 | 0.71 |
| 8.890 | 2-Methyl-1-undecanol | C<sub>12</sub>H<sub>26</sub>O | 186 | 4.61 |
| 10.256 | 4-Methylundecane | C<sub>12</sub>H<sub>26</sub> | 170 | 1.12 |
| 10.464 | 2,6,11-Trimethyldodecane | C<sub>15</sub>H<sub>32</sub> | 212 | 1.52 |
| 10.848 | 2-Methyloctadecane | C<sub>19</sub>H<sub>40</sub> | 268 | 1.27 |
| 12.122 | Oxalic acid, 6-ethyloct-3-yl isohexyl ester | C<sub>18</sub>H<sub>34</sub>O<sub>4</sub> | 314 | 0.57 |
| 12.481 | Sulfurous acid, pentadecyl 2-propyl ester | C<sub>18</sub>H<sub>38</sub>O<sub>3</sub>S | 334 | 2.77 |
| 12.822 | Tetradecanoic acid | C<sub>14</sub>H<sub>28</sub>O<sub>2</sub> | 228 | 3.30 |
| 17.992 | 2-Monopalmitin | C<sub>19</sub>H<sub>38</sub>O<sub>4</sub> | 330 | 3.33 |
| 30.806 | 3β-9,19-Cyclolanost-24-en-3-ol, acetate, | C<sub>32</sub>H<sub>52</sub>O<sub>2</sub> | 468 | 20.38 |
| 31.901 | 3β, 4α, 5α-4,14-Dimethyl-9,19-cycloergost-24(28)-en-3-ol acetate | C<sub>32</sub>H<sub>52</sub>O<sub>2</sub> | 468 | 25.40 |

**Table 4:** Compounds identified in *A. africana* extract known to serve as reducing agents, capping agents or both in nanoparticle synthesis.

| Compounds | Chemical Class | Reducing/Capping agent | References |
| --- | --- | --- | --- |
| Oxalic acid, 6-ethyloct-3-yl isohexyl ester | Ester | Reducing agent | Khandel *et al.* (2018) |
| Sulfurous acid, pentadecyl 2-propyl ester | Ester | Reducing agent | Khandel *et al.* (2018) |
| Tetradecanoic acid | Fatty acids | Reducing and capping agent | Afonso *et al.* (2024) |
| 2-Monopalmitin | Fatty acids | Reducing and capping agent | Afonso *et al.* (2024) |
| 3β-9,19-Cyclolanost-24-en-3-ol, acetate | Terpenoid | Reducing and capping agents | Mashwani *et al.* (2016) |
| 4,14-Dimethyl-9,19-cycloergost-24(28)-en-3-yl acetate | Terpenoid | Reducing and capping agents | Mashwani *et al.* (2016) |

**Table 5:** Compounds present on *A. africana*-derived nanoparticles.

| Nanoparticles | Retention Time (Min) | Name of compound | Molecular formula | Molecular weight (g/mol) | Peak Area % |
| --- | --- | --- | --- | --- | --- |
| AuNPs | 33.265 | 2-Monopalmitin | C<sub>19</sub>H<sub>38</sub>O<sub>4</sub> | 330 | 57.25 |
| AgNPs | 29.009 | 3,7-Dimethyloct-6-enyl isobutyl carbonate | C<sub>15</sub>H<sub>28</sub>O<sub>3</sub> | 256 | 7.66 |
|  | 33.665 | Dihydrosqualene | C<sub>30</sub>H<sub>52</sub> | 412 | 9.91 |
| CuO NPs | 20.166 | Octadecanal | C<sub>18</sub>H<sub>36</sub>O | 268 | 4.50 |

**Table 6:**
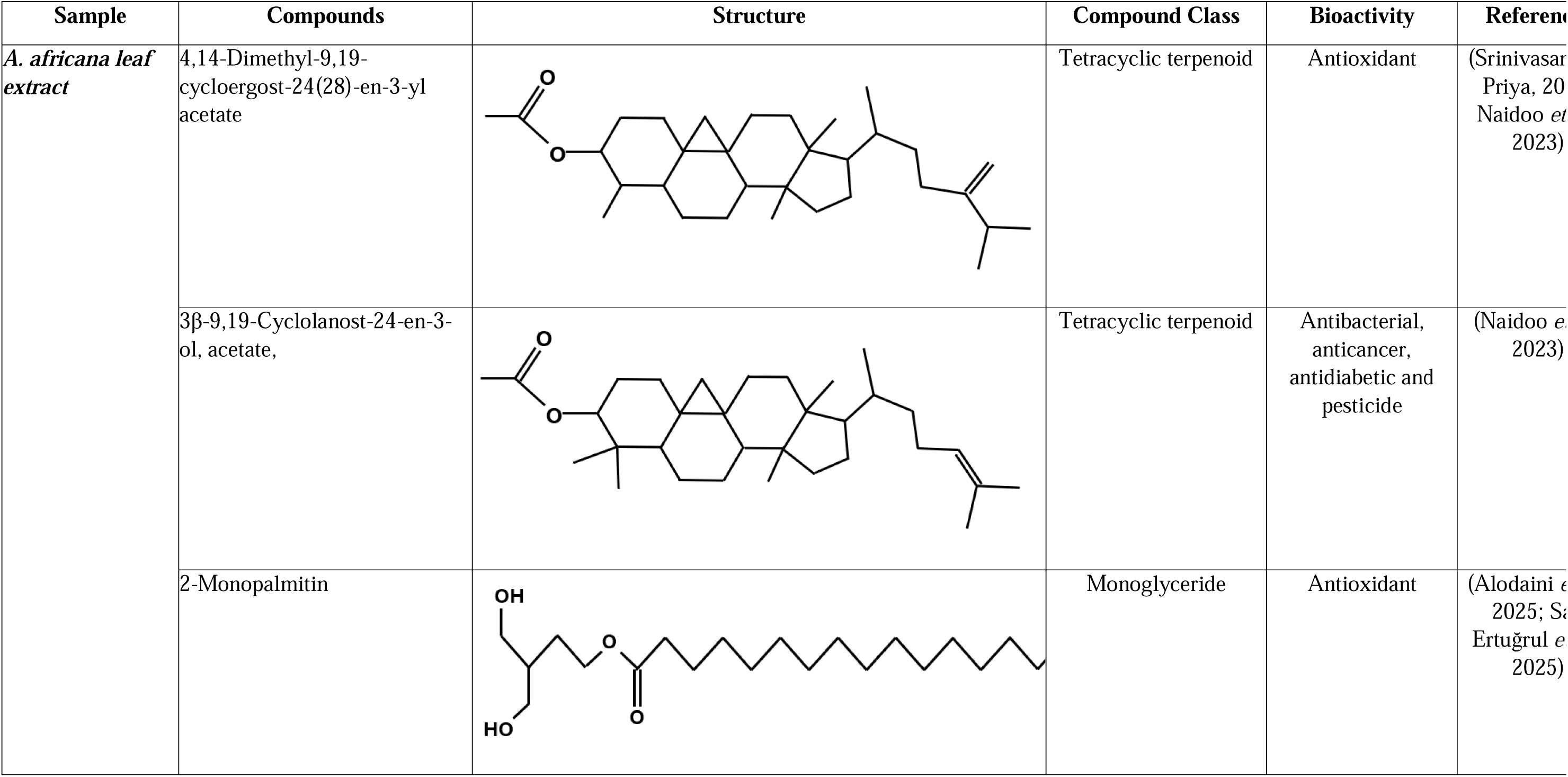
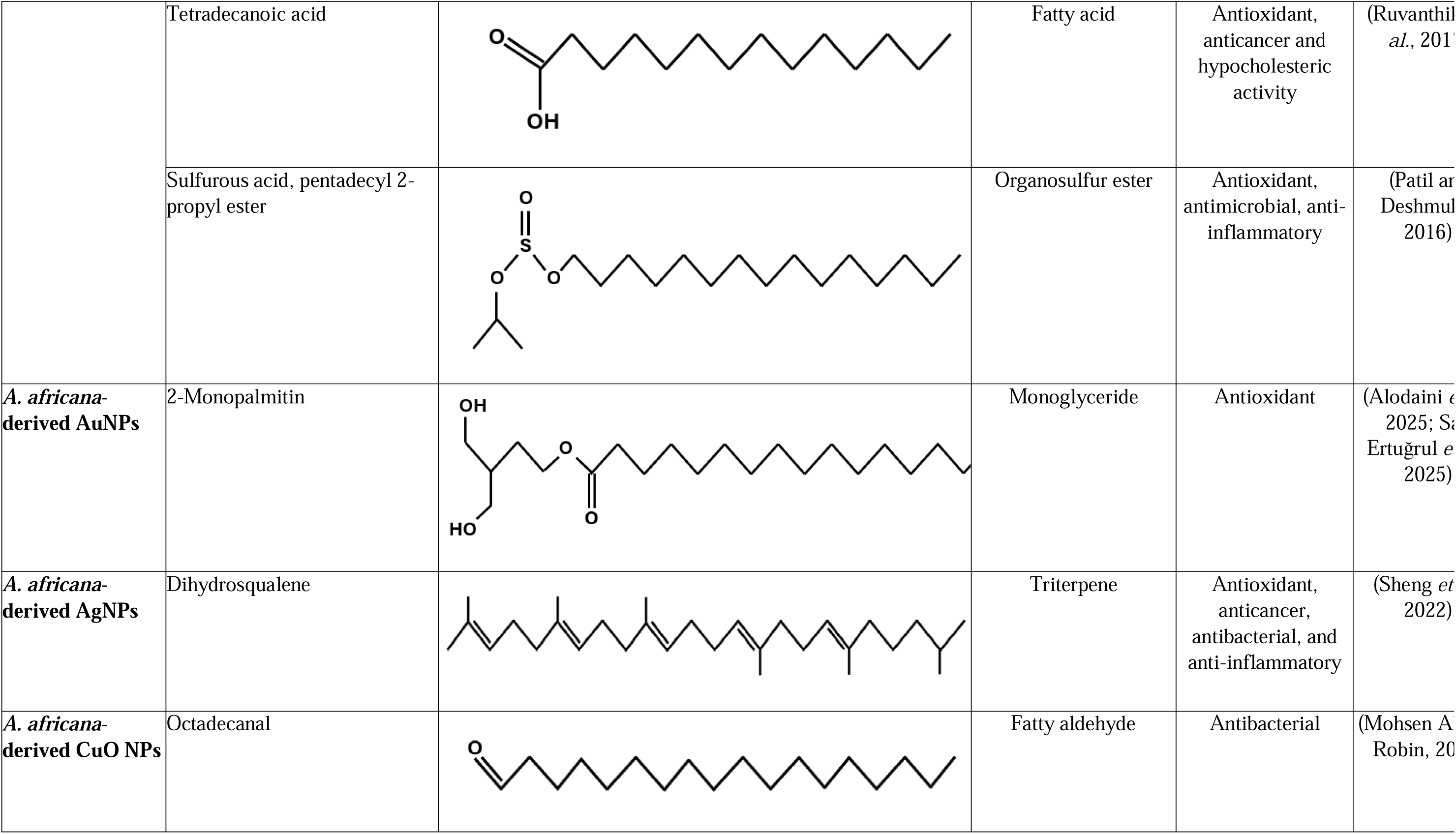
Structure and biological activity of compounds identified in *A. africana* leaf extracts and *A. africana*-derived nanoparticles.

GC-MS analysis of *A. africana* leaf extract identified a diverse range of bioactive compounds, including esters, terpenoids, fatty acids, monoglycerides and organosulfur compounds. These compounds contain functional groups such as hydroxyl (-OH), carbonyl (C=O), esters and alkyl chains. The AuNPs retained 2-monopalmitin from the plant extract on their surface, whereas the AgNPs and CuO NPs were coated with distinct compound, dihydrosqualene and octadecanal, that were not detected in the extract.

### 3.5. Antibacterial activity analysis

Various concentrations (6.25 – 200 µg/ml) of *A. africana* leaf extracts, and nanoparticles synthesised using *A. africana* leaf extracts and *M. magnetotacticum* were investigated against both Gram-negative and Gram-positive bacteria to evaluate their broad-spectrum antibacterial activity. Two Gram-negative bacteria, *Escherichia coli* and *Pseudomonas aeruginosa*, and two Gram-positive bacteria, *Enterococcus faecalis* and *Staphylococcus aureus* were used for evaluating antibacterial activity. The antibacterial activity Figure 4 illustrates the antibacterial activity of *A. africana* extracts, AuNPs, AgNPs and CuO NPs synthesised from *A. africana* and Figure 5 depicts the antibacterial activity of the nanoparticles derived from *M. magnetotacticum*. The results indicate the inhibition of bacterial growth relative to the growth of non-treated bacteria (uninhibited growth or negative control). Neomycin was used as a positive control. *A. africana*-derived AgNPs exhibited a dose-dependent inhibition across all tested bacteria, with increased inhibition at higher concentrations. These nanoparticles achieved up to 86% growth inhibition, with the highest growth inhibition being achieved at AgNP concentrations between 100 and 200 µg/mL. In contrast, *A. africana* extracts and *A. africana* AuNPs exhibited low antibacterial effects, displaying up to only 30% inhibition across all species tested. Similarly, *A. africana* CuO NPs also exhibited low antibacterial activity, however, moderate antibacterial activity with approximately 47% was observed against *E. coli* at concentrations between 50 and 200 µg/mL. The AuNPs, AgNPs and CuO NPs synthesised using *M. magnetotacticum* exhibited low antibacterial activity against all the tested bacterial species, displaying less than 25% inhibition. Overall, the plant synthesised nanoparticles displayed better bacterial growth inhibition compared to the bacterially synthesised nanoparticles. Among the *A. africana* nanoparticles, the AgNPs showed the best activity compared to the other metallic nanoparticles.

**Figure 4:**
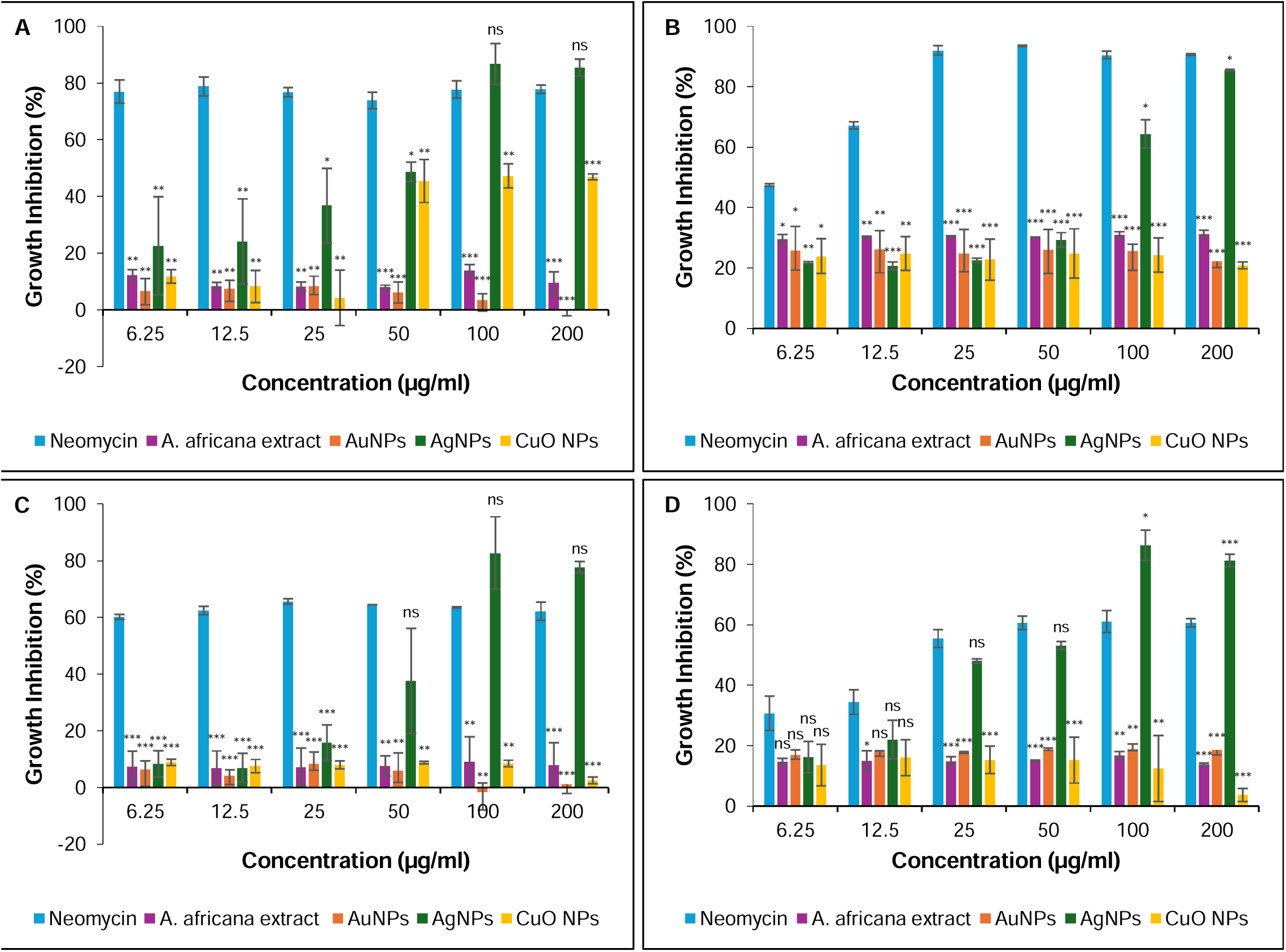
Antibacterial activity of *A. africana* leaf extract and AuNPs, AgNPs and CuO NPs synthesized from *A. africana* against (A) *E. coli*, (B) *E. faecalis*, (C) *P. aeruginosa* and (D) *S. aureus* after incubation for 24 hours at 30°C. Neomycin was used as a positive control. Bars represent mean ± STDEV. Statistical significance was evaluated using one-way ANOVA, * p < 0.05, ** p < 0.01, ***p < 0.001 and ns = not significant, when the positive control is compared to each NP at the given concentration.

**Figure 5:**
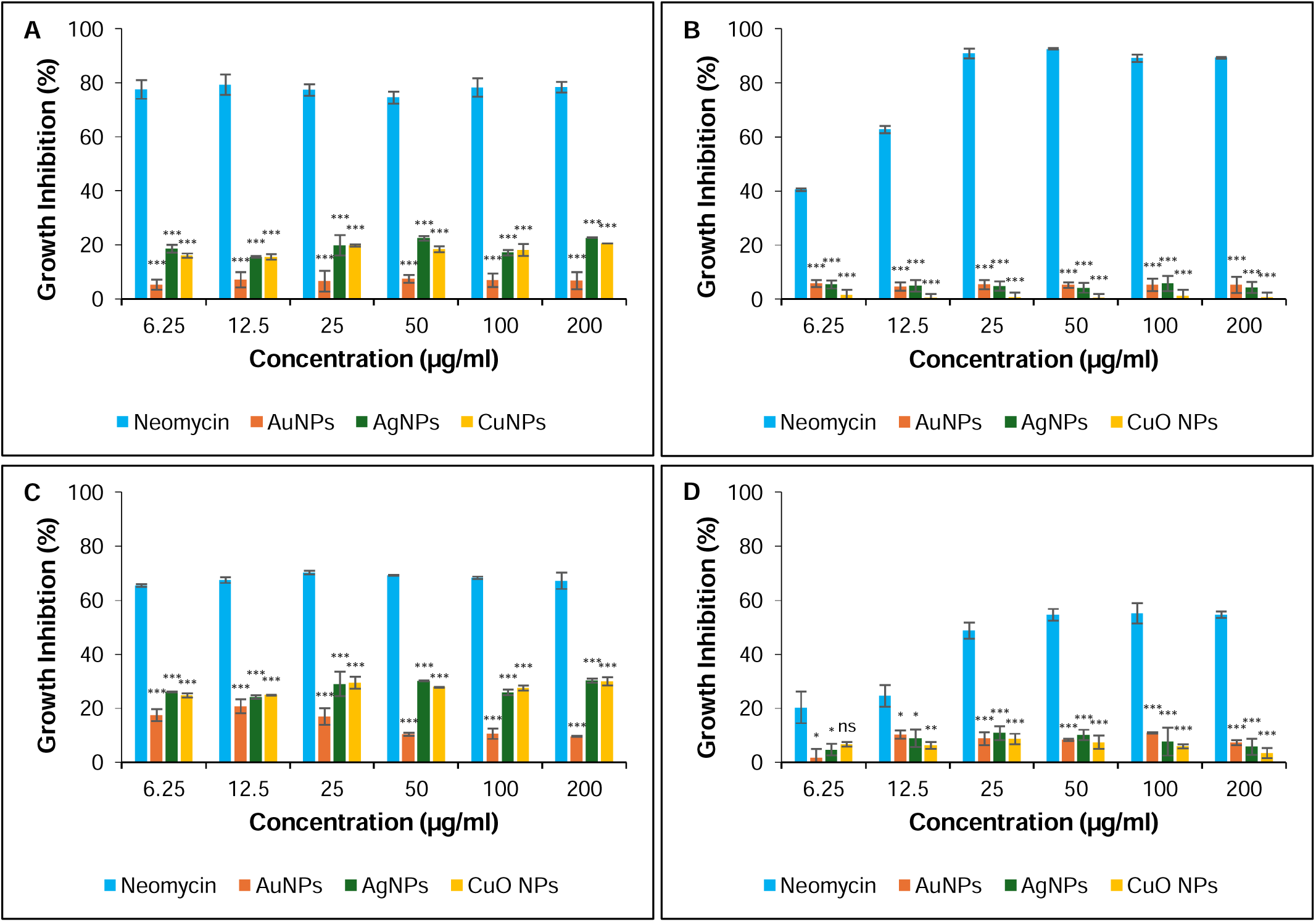
Antibacterial activity of AuNPs, AgNPs and CuO NPs synthesized from *M. magnetotacticum* against (A) *E. coli*, (B) *E. faecalis*, (C) *P. aeruginosa* and (D) *S. aureus* after incubation for 24 hours at 30°C. Neomycin was used as a positive control. Bars represent mean ± STDEV. Statistical significance was evaluated using one-way ANOVA, * p < 0.05, ** p < 0.01, ***p < 0.001 and ns = not significant, when the positive control is compared to each NP at the given concentration.

To assess the antibacterial potency of the *A. africana* AgNPs nanoparticles the minimum inhibitory concentrations (MIC) against the bacterial strains was determined. Table 7 compares the antibacterial potency of AgNPs derived from aloe leaf extracts to those synthesised in this study.

**Table 7:** Comparison of the MICs of AgNPs synthesized from various aloe leaf extracts against various bacteria.

| Plant species | Nanoparticle size (nm) | Nanoparticle Shape | Bacterial species | MIC ($\mu\text{g/ml}$) | References |
| --- | --- | --- | --- | --- | --- |
| *Aloe vera* | 9-18 | Spherical | *E. coli* | 780 | Yadav and Kumar (2016) |
|  |  |  | *E. faecalis* | 400 |  |
|  |  |  | *P. aeruginosa* | 780 |  |
|  |  |  | *S. aureus* | 195 |  |
| *Aloe vera* | 7-65 | Spherical | *E. coli* | 125 | Sharma *et al.* (2021) |
|  |  |  | *S. aureus* | 125 |  |
| *Aloe vera* | 15-25 | Spherical | *E. coli* | 8 | Ahmad *et al.* (2022) |
|  |  |  | *S. aureus* | 7.8 |  |
| *Aloe vera* | 30-80 | Spherical | *E. coli* | 50 | Arshad *et al.* (2022) |
|  |  |  | *P. aeruginosa* | 50 |  |
|  |  |  | *S. aureus* | 100 |  |
| *Aloe maculata* | 35.26 | Spherical | *E. coli* | 100 | Franceschinis *et al.* (2023) |
|  |  |  | *P. aeruginosa* | 25 |  |
|  |  |  | *S. aureus* | 200 |  |
|  | 55.38 | Spherical | *E. coli* | 12.5 |  |
|  |  |  | *P. aeruginosa* | 25 |  |
|  |  |  | *S. aureus* | 100 |  |
| *Aloe africana* | 11-30 | Spherical | *E. coli* | 100 | This study |
|  |  |  | *E. faecalis* | 200 |  |
|  |  |  | *P. aeruginosa* | 100 |  |
|  |  |  | *S. aureus* | 100 |  |

### 3.6. Cytotoxicity analysis

Since the *A. africana* leaf extract and the nanoparticles derived from this plant species exhibited antibacterial activity, their cytotoxicity was investigated at varying concentrations (31.25 – 1000 µg/ml) using the MTT assay. Figure 6 illustrates the cytotoxicity of the *A. africana* leaf extract AuNPs, AgNPs and CuO NPs synthesised from *A. africana* against HEK293 and HeLa cell lines. The results indicate cell survival relative to non-treated cells (negative control). Cisplatin was used as a positive control. *A. africana* extracts did not show any cytotoxic effects to the HEK293 and HeLa cell lines. Both AuNPs and CuO NPs displayed low cytotoxicity toward both cell lines across all tested nanoparticle concentrations. HEK293 cells treated with AuNPs exhibited up to 81.1% cell survival, while cells exposed to CuO NPs exhibited up to 87.7% cell survival. In HeLa cells, both AuNPs and CuO NPs consistently exhibited low cytotoxicity across all concentrations, with up to 80.5% and 87.8% cell survival, respectively. In contrast, AgNPs displayed a dose-dependent cytotoxic effect. The cytotoxicity of AgNPs towards HeLa cells mirrored that observed in HEK293 cells, displaying high cell survival at lower concentrations (31.25-125 µg/mL) and reduced cell survival at higher concentrations (250-1000 µg/mL). Overall, these finding show that *A. africana* AuNPs and CuO NPs are relatively non-cytotoxic to both HEK293 and HeLa cells, whereas AgNPs exhibit concentration-dependent cytotoxic effects in both cell lines. The AgNPs are cytotoxic at concentrations between 125-1000 µg/mL, whilst minimal toxicity is observed at 31.25 and 62.5 µg/mL.

**Figure 6:**
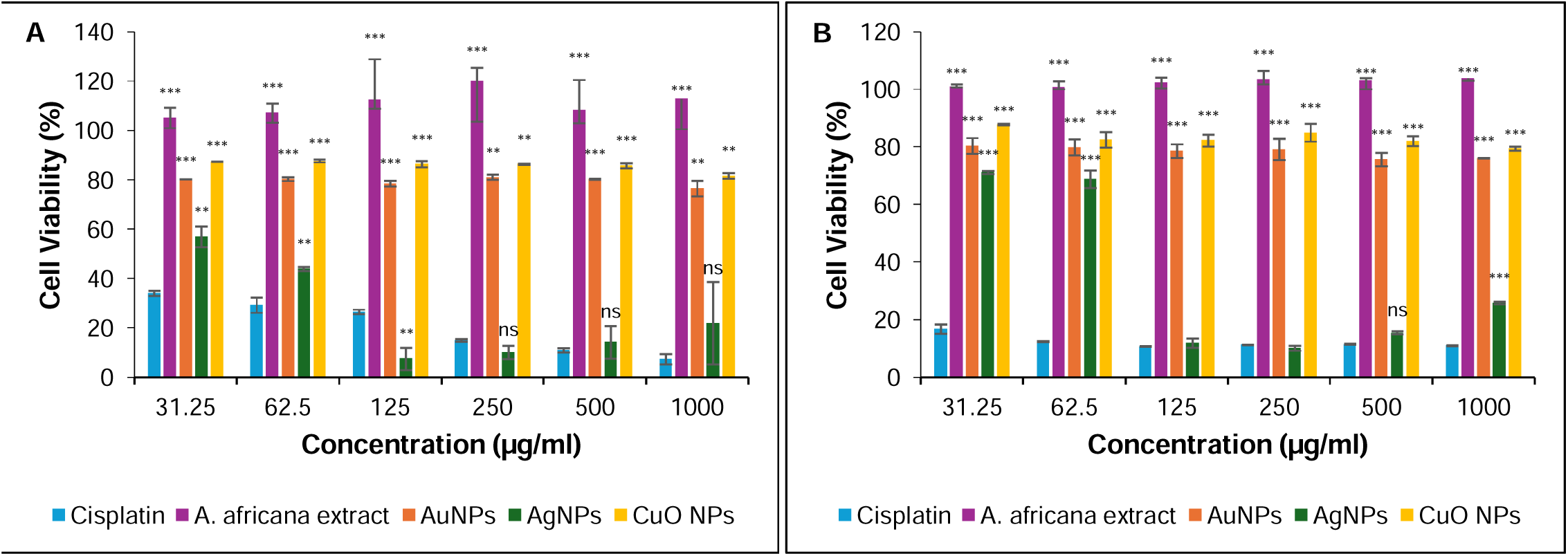
Cytotoxicity of cisplatin, *A. africana* extracts and *A. africana*-derived AuNPs, AgNPs and CuO NPs against (A) HEK293 and (B) HeLa cell lines. Cisplatin was used as a positive control. Bars represent mean ± sd. Statistical significance was evaluated using one-way ANOVA, * p < 0.05, ** p < 0.01, ***p < 0.001 and ns = not significant, when the positive control is compared to each NP at the given concentration.

## 4. Discussion

Green synthesis methods for nanoparticle production are non-toxic, pollution free, environmentally friendly and cost-effective, making them a more suitable alternative to conventional chemical and physical synthesis methods (Nadaroglu *et al*., 2017; Ying *et al*., 2022). In the present study, two green synthesis methods for nanoparticle synthesis were compared for preparing metallic nanoparticles. These involved the use of aqueous leaf extracts of *A. africana* and the magnetotactic bacteria, *M. magnetotacticum*, for the reduction of gold chloride, silver nitrate and copper nitrate solutions. UV-vis analysis confirmed the formation of the metallic nanoparticles using *A. africana*. It has been reported in literature that metallic nanoparticles show characteristic peaks at ranges of 500-600 nm for AuNPs, 400-500 nm for AgNPs and 250-350 nm for CuO NPs (Ashraf *et al*., 2016; Elbagory *et al*., 2016; Manjari *et al*., 2017). The nanoparticles synthesised in this study fall within these ranges, thereby verifying the presence of the nanoparticles of interest. UV-vis spectral analysis of nanoparticles synthesized using *M. magnetoatcticum* is not reported in this study, however, TEM-EDX analysis was used for primary confirmation of nanoparticle synthesis from these bacterial species as shown by Sancho *et al*. (2023) and Murei *et al*. (2021).

The size distributions and morphological diversity of *A. africana*- and *M. magnetotacticum*-derived AuNPs, AgNPs, and CuO NPs closely match with those reported for other Aloe species and bacterial systems. In plant-mediated synthesis, variability in morphology, displayed by the *A. africana* AuNPs likely arises from phytochemicals in plant extracts and reaction conditions used for synthesis, all of which critically influence nanoparticle size and shape (Hanna *et al*., 2025). Notably, *A. africana* AuNPs from this study differ from those synthesised by Machine *et al*. (2025) using the same plant, highlighting how synthesis parameters such as pH, temperature, metal precursor concentration, and reaction duration affects nucleation and growth processes resulting in distinct nanoparticle morphologies (Kazemi *et al*., 2023). Similarly, AuNPs synthesised by *M. magnetotacticum* in the study by Murei *et al*. (2021) showed comparable sizes but different shapes in comparison to this study. While species-specific patterns emerge in nanoparticle synthesis among bacteria, similarities in nanoparticle size and shape occur between bacterial species such as *Streptomyces albogriseolus*, *Bacillus* sp. and *Serratia* sp. ZTB29 are seen (Vithiya *et al*., 2014; Singh *et al*., 2023; El-Naggar *et al*., 2024).

Nanoparticle stability is crucial for functional performance in applications like biomedicine, environmental science, and engineering where aggregation compromises nanoparticle function (Femina and Kamalesh, 2024). In this study, the nanoparticles exhibited colloidal stability ranging from relatively stable to highly stable. For *A. africana*-derived nanoparticles, terpenoid and monoglyceride capping agents provide steric hindrance and electrostatic repulsion to prevent aggregation and maintain functionality (Javed *et al*., 2022). *M. magnetotacticum* nanoparticles owe their stability to encapsulation by the magnetosome membrane, which acts as a natural capping layer (Javed *et al*., 2022).

Plant-mediated nanoparticle synthesis also relies heavily on the phytochemicals in the extract, serving as reducing and capping agents. Diverse bioactive compounds were identified in the *A. africana* leaf extract including esters, terpenoids, monoglycerides and fatty acids. These compounds contain functional groups such as hydroxyl (-OH), carbonyl (C=O), esters and alkyl chains which facilitate the reduction and stabilisation of nanoparticles (Khandel *et al*., 2018). Interestingly, only the AuNPs retained the compound 2-monopalmitin from the parent *A. africana* leaf extract on their surface, while AgNPs and CuO NPs were coated with distinct compounds not detected in the extract. The presence of compounds on nanoparticles that were absent from the parent extract is a known phenomenon in green synthesis. This occurs when diverse phytochemicals undergo oxidation, rearrangement or hydrolysis during metal ion reduction forming new compounds that coat nanoparticle surfaces (Amini and Akbari, 2019; Singh *et al*., 2023; Rana *et al*., 2025).

While previous green synthesis studies relied primarily on FTIR for nanoparticle surface chemistry, our sonication-GC-MS approach uniquely identified desorbed capping agents specifically 2-monopalmitin on AuNPs, dihydrosqualene on AgNPs and octadecanal on CuO NPs which are phytochemicals that are known to enhance colloidal stability and bioactivity. In a previous study by Raut *et al*. (2014) FTIR’s limitations were illustrated by the analysis of *Withania somnifera* AgNPs which suggested capping by sitoindoside. However, further analysis using HPLC confirmed catechin, p-coumaric acid, luteolin-7-glucoside, and withanolide derivatives instead demonstrating FTIR’s functional group inference versus chromatography’s compound-specific identification, highlighting potential discrepancies in assigning reduction/capping agents during green synthesis.

Octadecanal detected on CuO NPs likely arose from the reduction/decarboxylation of the fatty acid precursors tetradecanoic acid and 2-monopalmitin identified from the plant extract, analogous to established fatty acid-to-aldehyde conversion mechanisms (Akhtar *et al*., 2013). Dihydrosqualene detected on AgNPs could be a product from carbocation-mediated ring opening of the tetracyclic terpenoids cyclolanostanes and ergostanes from the extract, a mechanism characteristic of terpenoid cyclase cascades (Christianson, 2017). Interestingly, this conversion to dihydrosqualene displays a reverse of the natural squalene to cyclic terpenoid conversion (Christianson, 2017). Importantly, many of these compounds identified in the *A. africana* extract and nanoparticles are reported to possess antibacterial activity. Despite the antibacterial activity of these compounds, the extract alone exhibited limited antibacterial activity. However, when combined with metallic cores to form nanoparticles, particularly for AgNPs, a synergistic effect is shown displaying improved antibacterial activity. This also points to the mechanism of action of AgNPs which is influenced by the physicochemical properties of the nanoparticles such as their size, shape, stability and composition.

The *A. africana* extracts displayed minimal antibacterial activity against *E. coli*, *E. faecalis*, *P. aeruginosa*, and *S. aureus*, although compounds with antibacterial activity were identified from these extracts from GC-MS analysis. Nanoparticle formation enhanced potency especially for *A. africana* AgNPs, achieving dose-dependent inhibition up to 86% against all strains showing broad spectrum activity. These nanoparticles outperformed the *A. africana* AuNPs and CuO NPs. The minimum inhibitory concentrations (MICs) of *A. africana*-derived AgNPs are comparable to those from other aloe species reported by Sharma *et al*. (2021) and (Franceschinis *et al*., 2023) For instance, *A. vera*-derived AgNPs showed similar activity against *E. coli* and *S. aureus* (Sharma *et al*., 2021), while some larger particles achieved lower MICs (Ahmad *et al*., 2022). Despite this, the smaller *A. africana*-derived AgNPs in this study achieved potency at moderate concentrations, highlighting plant-specific phytochemical capping as key to their efficacy. The small size and spherical shape of the AgNPs enabled interaction with the cell membrane, penetration and subsequently disruption of crucial cellular processes leading to effective antibacterial action (Ali *et al*., 2024; Liknaw *et al*., 2025). The AuNPs and CuO NPs were 4-10 times larger in size and their diverse and irregular shapes could have limited their interaction with the bacterial membrane. Additionally, AgNPs release Ag^+^ ions that alter phospholipid bilayers/cell walls, increase permeability, disrupt electron transport, and target intracellular DNA and enzymes (Girma, 2023; Jiang *et al*., 2024).

In contrast, *M. magnetotacticum*-derived nanoparticles exhibited poor bacterial growth inhibition despite being small and spherical in shape. This indicated that in this instance size and shape did not influence their antibacterial effectiveness but rather the surface capping agents played a crucial role in their antibacterial activity (Franceschinis *et al*., 2023). It has previously been shown that *M. magnetotacticum*-derived nanoparticles were surrounded by a phospholipid bilayer membrane containing proteins that lacked antibacterial effects (Yan *et al*., 2017). The importance of the surface capping agents was further demonstrated by Murei *et al*. (2021) who showed that *Pyrenacantha grandiflora* extract conjugation to *M. magnetotacticum* AuNPs exhibited antibacterial activity unlike unconjugated *M. magnetotacticum* AuNPs. This suggests that the *P. grandiflora* extract capped the AuNPs with antibacterial phytochemicals that influenced the antibacterial effects of these nanoparticles.

Overall, the *A. africana* AgNPs outperformed *A. africana*-derived AuNPs and CuO NPs and *M. magnetotacticum*-derived AuNPs, AgNPs and CuO NPs using a dual antibacterial mechanism i.e. by possessing antibacterial phytochemicals and using the metallic mechanism of action of AgNPs, unlike single-target antibiotics. The mechanism of action of traditional antibiotics is relatively simple, which partially explains the rapid emergence of bacterial resistance (Wang *et al*., 2017). Antibiotics are classified according to their mechanism of action, grouping them by how they work (Uddin *et al*., 2021). These mechanisms include inhibiting the synthesis of the cell wall or inhibiting the synthesis of proteins or nucleic acids or metabolites or disrupting metabolic pathways or the cell membrane (Kapoor *et al*., 2017; Uddin *et al*., 2021). In contrast, nanoparticles exert bactericidal effects through multiple modes of action as described above (Wang *et al*., 2017). This multifaceted approach is important for their sustained antibacterial activity, and it is unlikely that bacteria will harbour multiple mutated genes necessary to resist all mechanisms of action of nanoparticles, making it difficult for them to develop resistance to nanoparticles (Wang *et al*., 2017).

The *A. africana* AgNPs show immense potential for use as antibacterial agents and most importantly present a viable alternative to antibiotics. However, the biocompatibility of nanoparticles is an essential component that ensures safe nanoparticle use in biomedical applications (Kyriakides *et al*., 2021; Eyube, 2024). In this study it was observed that although *A. africana* AuNPs and CuO NPs showed limited antibacterial activity, they were found to be non-cytotoxic towards both normal and cancerous cells. Similarly, Machine *et al*. (2025) reported that *A. africana* AuNPs were biocompatible with up to 90% viability when in contact with cancer cell lines (Machine *et al*., 2025). CuO NPs synthesized from *Athrixia phylicoides* exhibited no cytotoxicity toward HEK293 cells, aligning with the non-cytotoxic profile of the CuO NPs in this study (Kaningini *et al*., 2023).

The cytotoxicity of *A. africana* AgNPs although displaying the best antibacterial activity was found to be dose dependent i.e. higher concentrations of these nanoparticles were cytotoxic. Similarly, AgNPs derived from *Rhizophora apiculata* leaf extract displayed up to 100% cell death in HEK293 and up to 75% cell death in HeLa cells (Liu *et al*., 2021). Likewise, AgNPs synthesised from *Phlogacanthus thyrsiformis* effectively inhibited HeLa cells (Kumar *et al*., 2017). Factors affecting the cytotoxicity of the AgNPs are similar to those affecting their antibacterial activity i.e. the size, shape and composition. Smaller sized nanoparticles are more cytotoxic due to their ability to penetrate cell membranes, resulting in cell disruption (Kumar *et al*., 2025). The *A. africana* AgNPs had the smallest size compared to the AuNPs and CuO NPs and therefore were cytotoxic. These findings were consistent with previously reported plant-derived nanoparticles, AgNPs exhibited dose-dependent cytotoxic effects (Kumar *et al*., 2017; Liu *et al*., 2021). However, AgNPs are known for their medicinal use for example in nano-hydrogels for wound healing, as the slow release of AgNPs mitigates AgNP cytotoxicity thereby enabling application in wound healing (Haidari *et al*., 2021). Another factor affecting the cytotoxicity of AgNPs is uncontrolled/rapid release and excessive accumulation of Ag^+^ ions, therefore hydrogels serve as a reservoir for controlled release of Ag^+^ maintaining antibacterial effects with minimal cytotoxicity (Haidari *et al*., 2021). The *A. africana* AgNPs showed cytotoxicity at 125-1000 µg/mL against both cell lines which overlaps with the peak antibacterial activity range of 100-200 µg/mL. However, hydrogel formulations present a possible solution for addressing this. Therefore, the *A. africana* AgNPs in this study show potential for incorporation into a hydrogel for wound healing.

## 5. Conclusion

This study provides an in-depth characterisation of AuNPs, AgNPs and CuO NPs synthesised using *A. africana* leaf extract and *M. magnetotacticum*, with focus on their physicochemical properties, antibacterial and cytotoxic activities. Characteristic peaks confirmed the successful synthesis of the respective nanoparticles. *A. africana-*mediated synthesis of metallic nanoparticles led to the synthesis of nanoparticles displaying diverse morphologies and size distributions. This diversity is likely due to the phytochemical and precursor variations which influence nanoparticle formation. *M. magnetotacticum* favoured formation of spherical nanoparticles with size distributions comparable to what is reported in literature, although some nanoparticles had distinct features and broader size ranges. EDX analysis confirmed the elemental composition of the biosynthesised nanoparticles confirming the presence of the respective metals. All the nanoparticles showed moderate dispersity with PDI values between 0.25 and 0.4 and zeta potentials of up to -36.7 mV, suggesting moderate dispersity and moderate to good colloidal stability. GC-MS analysis of *A. africana* extract identified terpenoids, monoglycerides, esters, and fatty acids that served as reducing and capping agents. Notably, AuNPs retained 2-monopalmitin from the extract, while AgNPs and CuO NPs featured transformed compounds (dihydrosqualene, octadecanal) formed during synthesis, enhancing nanoparticle functionality and stability.

*A. africana*-derived AgNPs displayed good antibacterial efficacy against both Gram-negative and Gram-positive bacteria, with complete inhibition observed at higher concentrations, making them excellent candidates for further evaluation. Although the *A. africana*-derived AuNPs and CuO NPs demonstrated limited antibacterial effects, they exhibited low cytotoxicity toward HEK293 and HeLa cell lines, with over 80% survival across the tested concentrations. In contrast, AgNPs showed concentration-dependent toxicity, with reduced and enhanced cell survival at certain concentrations, highlighting the need for controlled-release formulations like hydrogels. *M. magnetotacticum*-derived nanoparticles showed limited antibacterial activity across all strains despite favourable size and shape morphologies which may have been attributed to the non-antibacterial magnetosome membrane capping.

Overall, *A. africana* AgNPs demonstrate significant broad-spectrum antibacterial potential with defined phytochemical synergies, making them promising candidates for hydrogel-based wound healing applications.

## Data accessibility

The University of KwaZulu-Natal has a central repository, Yabelana to archive all research data, raw or refined. The link to the website is https://yabelana.ukzn.ac.za/.

## Author contributions

AG and KP conceived and designed the project, KN, AG and NN acquired the data, KN, AG, KP and NN analysed and interpreted the data, KN wrote the paper, KN, AG, KP and NN edited the paper.

## Acknowledgements

This work was funded by the National Research Foundation of South Africa (Grant Number: SRUG2203291132). The funding body, however, had no role in study design; in the collection, analysis and interpretation of data; in the writing of the report; and in the decision to submit the data for conference presentation.

## Abbreviations

AuNPs: Gold nanoparticles
AgNPs: Silver nanoparticles
CuO NPs: Copper oxide nanoparticles
MTB: Magnetospirillum magnetotacticum

