## Supplementary material for "Comparison of the bioactivity of gold, silver and copper oxide nanoparticles derived from *Aloe africana* leaf extract and *Magnetospirillum magnetotacticum*": Table S1

**Supplementary Information**

**Table S1.** Reported studies on the compounds identified in the *A. africana* extracts and nanoparticles in this study.

| **Sample** | **Compound** | **Plant species** | **Reference** |
| --- | --- | --- | --- |
| *Aloe africana* extract | 4,8-Dimethylundecane | *Zanthoxylum myriacanthum* | Dai *et al.* (2023) |
|  | 2-Hexadecen-1-ol | *Schimpera arabica* | Hidayathulla *et al.* (2018) |
|  | 2-Methyl-1-undecanol | *Foeniculum vulgare* | (Gatea, 2025) |
|  | 4-Methylundecane | *Artemisia argyi* | (Gu *et al.*, 2024) |
|  | 2,6,11-Trimethyldodecane | *Zanthoxylum myriacanthum* | (Dai *et al.*, 2023) |
|  | 2-Methyloctadecane | *Solenostemma arghel* | (Abdel-Motaal *et al.*, 2022) |
|  | Oxalic acid, 6-ethyloct-3-yl isohexyl ester | *Anogeissus pendula* | (Bairwa *et al.*, 2018) |
|  | Sulfurous acid, pentadecyl 2-propyl ester | *Psychotria dalzellii* | (Thejashree and Naika, 2024) |
|  | (3.beta.)-9,19-Cyclolanost-24-en-3-ol, acetate | *Cassia angustifolia* | (Al-Marzoqi *et al.*, 2016) |
|  | Palmitic acid/N-Hexadecanoic Acid | *Aloe aculeata*  *Aloe africana*  *Aloe arborescens*  *Aloe barbadensis*  *Aloe ferox*  *Aloe marlothii*  *Aloe spectabilis*  *Aloe vera* | (Andrea *et al.*, 2020) (Kushwaha *et al.*, 2023) |
|  | 2-Monopalmitin | *Olea europea L.* | (Vural and Akay, 2021) |
|  | (3.beta.,4.alpha.,5.alpha.)-9,19-Cycloergost-24(28)-en-3-ol, 4,14-dimethyl-, acetate, | *Cenchrus ciliaris*  *Cenchrus setigerus* | (Singariya *et al.*, 2012) |
|  | Tetradecanoic acid/ myristic acid | *Aloe aculeata*  *Aloe africana*  *Aloe arborescens*  *Aloe barbadensis*  *Aloe ferox*  *Aloe marlothii*  *Aloe spectabilis*  *Aloe vera* | (Andrea *et al.*, 2020) (Kushwaha *et al.*, 2023) |
| *A. africana* AuNPs | 2-Monopalmitin | *Olea europea L.* | (Vural and Akay, 2021) |
| *A. africana* AgNPs | 3,7-Dimethyloct-6-enyl isobutyl carbonate | *Physalis minima Linn.* | (Sowmiya *et al.*, 2021) |
|  | Dihydrosqualene | *Anisomelis indica* | (Deshmukh, 2024) |
| *A. africana* CuO NPs | Octadecanol | *Aloe vera* | Kushwaha *et al.* (2023) |

Abdel-Motaal, F. F., Maher, Z. M., Ibrahim, S. F., El-Mleeh, A., Behery, M., & Metwally, A. A. (2022). Comparative Studies on the Antioxidant, Antifungal, and Wound Healing Activities of *Solenostemma arghel* Ethyl Acetate and Methanolic Extracts. *Applied Sciences*, *12*, 1-22

Al-Marzoqi, A., Hadi, M., & Hameed, I. (2016). Determination of metabolites products by *Cassia angustifolia* and evaluate antimicobial activity. *Journal of Pharmacognosy and Phytotherapy*, *8*, 25-48

Andrea, B., Dumitrița, R., Florina, C., Francisc, D., Anastasia, V., Socaci, S., & Adela, P. (2020). Comparative analysis of some bioactive compounds in leaves of different Aloe species. *BMC Chem*, *14*, 1-11

Bairwa, S., saini, s., Sharma, M., & Agrawal, R. D. (2018). Isolation and identification of phytosterols from Anogeissus pendula (Edgew) and their antimicrobial potency. *Journal of Pharmacognosy and Phytochemistry*, *8*, 1665-1670

Dai, W., Zhang, L., Dai, L., Tian, Y., Ye, X., Wang, S., Li, J., & Wang, Q. (2023). Comparative Analysis of Chemical Composition of *Zanthoxylum myriacanthum* Branches and Leaves by GC-MS and UPLC-Q-Orbitrap HRMS, and Evaluation of Their Antioxidant Activities. *Molecules*, *28*, 1-15

Deshmukh, D. (2024). GC-MS analysis and conservation of ethnomedicinal aromatic plant Anisomelis indica (L). *International Journal of Pharmacognosy and Life Science*, *5*, 53-56

Gatea, A. (2025). Metabolite Profiling Using FTIR and GC-MS Techniques and Bioactivities of Fennel (*Foeniculum vulgare*) Flowers a Traditional Iraqi Medicinal Plant. *South Asian Research Journal of Pharmaceutical Sciences*, *7*, 70-76

Gu, H., Peng, Z., Kuang, X., Hou, L., Peng, X., Song, M., & Liu, J. (2024). Enhanced Synthesis of Volatile Compounds by UV-B Irradiation in *Artemisia argyi* Leaves. *Metabolites*, *14*, 1-14

Hidayathulla, S., Shahat, A. A., Ahamad, S. R., Al Moqbil, A. A. N., Alsaid, M. S., & Divakar, D. D. (2018). GC/MS analysis and characterization of 2-Hexadecen-1-ol and beta sitosterol from *Schimpera arabica* extract for its bioactive potential as antioxidant and antimicrobial. *J Appl Microbiol*, *124*, 1082-1091

Kushwaha, P., Shrivastava, M., Pandey, D., Verma, K., & Dwivedi, L. (2023). The phytochemical profiling of *Aloe vera* through GC-MS and compounds activity validation at NCBI for industrial value addition of the plant. *South Asian Journal of Experimental Biology*, *13*, 20-32

Singariya, P., Mourya, K., & Kumar, P. (2012). *In vitro* Studies of Antimicrobial Activity of Crude Extracts of the Indian Grasses Dhaman (*Cenchrus ciliaris*) and Kala-Dhaman (*Cenchrus setigerus*). *Indian journal of pharmaceutical sciences*, *74*, 261-265

Sowmiya, V., Sakthi, P. S., & Kumar, P. (2021). GC-MS profiling of different fractions of *Physalis minima Linn.*, leaves. *Natural Volatiles &amp; Essential Oils*, *8*, 12852–12859

Thejashree, M., & Naika, R. (2024). Identification of Bioactive Compounds in Acetone Leaf and Stem-Bark Extracts of *Psychotria dalzellii* Hook.f. by GC-MS Analysis and Evaluation of in vitro Anti-bacterial Properties. *Asian Journal of Biological and Life Sciences*, *12*, 499-509

Vural, N., & Akay, M. (2021). Chemical compounds, antioxidant properties, and antimicrobial activity of olive leaves derived volatile oil in West Anatolia. *Journal of the Turkish Chemical Society Section A: Chemistry*, *8*, 511-518
